# Single-Cell Analysis Reveals Distinct Immune and Smooth Muscle Cell Populations that Contribute to Chronic Thromboembolic Pulmonary Hypertension

**DOI:** 10.1101/2022.01.26.477878

**Authors:** Gayathri Viswanathan, Hélène Fradin Kirshner, Nour Nazo, Asvin Ganapathi, Ian Cummings, Issac Choi, Anmol Warman, Susana Almeida-Peters, John Haney, David Corcoran, Yen-Rei Yu, Sudarshan Rajagopal

**Author notes:** To whom correspondence should be addressed: Yen-Rei Yu, MD, PhD, Division of Pulmonolary, Allergy, and Critical Care Medicine, Department of Medicine, Box, DUMC, Durham, NC 27710; Sudarshan Rajagopal, MD, PhD, Division of Cardiology, Department of Medicine, Box 102147, DUMC, Durham, NC 27710. Division of Cardiac Surgery, Department of Surgery, Ohio State Medical Center, Columbus, OH, USA. These authors contributed equally to this work (Yen-Rei Yu, Sudarshan Rajagopal).

## Abstract

Chronic thromboembolic pulmonary hypertension (CTEPH) is a sequelae of acute pulmonary embolism (PE) in which the PE remodels into a chronic scar in the pulmonary arteries. This results in vascular obstruction, small vessel arteriopathy and pulmonary hypertension. Our current understanding of CTEPH pathobiology is primarily derived from cell-based studies limited by the use of specific cell markers or phenotypic modulation in cell culture. Here we used single cell RNA sequencing (scRNAseq) of tissue removed at the time of pulmonary thromboendarterectomy (PTE) surgery to identify the multiple cell types, including macrophages, T cells, and smooth muscle cells, that comprise CTEPH thrombus. Notably, multiple macrophage subclusters were identified but broadly split into two categories, with the larger group characterized by an upregulation of inflammatory signaling predicted to promote pulmonary vascular remodeling. Both CD4+ and CD8+ T cells were identified and likely contribute to chronic inflammation in CTEPH. Smooth muscle cells were a heterogeneous population, with a cluster of myofibroblasts that express markers of fibrosis and are predicted to arise from other smooth muscle cell clusters based on pseudotime analysis. Additionally, cultured endothelial, smooth muscle and myofibroblast cells isolated from CTEPH thrombus have distinct phenotypes from control cells with regards to angiogenic potential and rates of proliferation and apoptosis. Lastly, our analysis identified protease-activated receptor 1 (PAR1) as a potential therapeutic target that links thrombosis to chronic PE in CTEPH, with PAR1 inhibition decreasing smooth muscle cell and myofibroblast proliferation and migration. These findings suggest a model for CTEPH similar to atherosclerosis, with chronic inflammation promoted by macrophages and T cells driving vascular remodeling through smooth muscle cell modulation, and suggest new approaches for pharmacologically targeting this disease.

## Introduction

CTEPH is defined as an elevation of pulmonary pressure and vascular resistance due to the presence of chronic pulmonary thrombi.^1^ In CTEPH, the acute PE does not dissolve in response to anticoagulants, but instead develops into an organized, scar-like, chronic “thrombus”, leading to persistent obstruction of the pulmonary vasculature.^2,3^ Approximately 1-2% of patients of acute PE go on to develop CTEPH, with an estimated incidence of over 10,000 patients in the United States annually.^1^ Treatment options for this highly morbid disease range from pulmonary thromboendarterectomy (PTE), a surgery to remove the thrombus only available to operable candidates (who have surgically accessible disease with no contraindications to surgery) to interventional therapies, such as balloon pulmonary angioplasty (BPA), and medical therapies that have only been shown to improve short-term endpoints.^4^ Thus, the development of novel medical therapies for CTEPH is currently a significant unmet need. However, this is currently limited by a knowledge gap in our understanding of CTEPH pathobiology.

The pathobiology of CTEPH differs significantly from acute PE. In acute PE, the fresh clots are red, easily detached from the pulmonary artery wall and consist mainly of red cells and platelets in a fibrin mesh.^5^ This is in contrast to CTEPH, where the chronic clots are yellow, incorporated in the pulmonary vascular wall and contain collagen, elastin and inflammatory cells.^3,5^ Numerous studies have focused on identifying potential pathogenic cells in CTEPH thrombus. These studies, while providing important insights into disease pathobiology, have only provided limited information on the heterogeneous cell populations that compose CTEPH thrombus.^6^ Some cells within chronic thrombus have been described as multipotent mesenchymal progrenitor cells^7^, myofibroblast-like cells, endothelial-like cells ^7-9^, or “sarcoma-like”^10,11^. Others have described the marked presence of inflammatory cells (B and T lymphocytes, macrophages and neutrophils)^12,13^. Due to the methodology used to examine the cells, it is unclear if these terms describe a single cell type or a heterogeneous cell population. In addition, two major limitations apply to the majority of these studies.^8-10,14,15^ First, is histopathology based on a set of limited markers that may have limited specificity or may be silent to relevant cell populations. Second, is a reliance on culturing cells from thrombus, thereby enriching for cells that grow under culture conditions that can result in phenotypic modulation.

Single cell technologies, most prominent among them single cell RNA sequencing (scRNAseq), have revolutionized the study of cardiovascular diseases, allowing the identification of rare cell types, the analysis of single-cell trajectory construction and the identification of novel drug targets^16^ Here we have used scRNAseq of samples of CTEPH thrombus removed at the time of PTE surgery to identify specific immune and mesenchymal cells in CTEPH thrombus that are predicted to promote vascular remodeling and inflammation. We also found that cells cultured from CTEPH thrombus displayed distinct phenotypes compared to control cells, a phenotype that could be modulated pharmacologically, suggesting an approach to develop novel CTEPH treatments.

## Materials and Methods

### Human specimens and patient clinical characteristics

Five patients undergoing pulmonary thromboendarterectomy surgery at Duke University Hospital were enrolled in an ongoing clinical study approved by the Duke University Medical Center Institutional Review Board (IRB number Pro00082338). Eligible subjects gave informed written consent prior to surgery. Exclusion criteria included age < 18 years old or inability to provide informed consent. Peripheral venous blood and PTE samples were obtained from each patient. **Supplementary Table 1** summarizes the demographic and clinical characteristics of the cohort. Preoperative imaging (CT angiogram of chest and VQ scan) for selected subjects is shown in **Supplementary Figure 1**.

### Sample collection and preparation

PTE surgery was performed at Duke University Hospital. Patients were placed on cardiopulmonary bypass and had the specimens removed under deep hypothermic circulatory arrest. After the endarterectomized material (“thrombus”) was removed, a portion of the proximal and distal portions of the thrombus were dissected and placed into sterile phosphate-buffered saline on ice. An on-call research team was notified and the samples were taken to the lab for further treatment. Thrombus removed at the time of surgery for all subjects included in this study is shown in **Supplementary Figure 2**. We first tested whether the proximal (main/lobar) or distal (segmental/subsegmental) were cellular. Flow cytometry demonstrated that the proximal portions of thrombus were fibrotic and largely acellular (data not shown), consistent with previous studies^17^. The portion of distal sample were separated for single cell RNA sequencing, paraffin embedding, OCT cryopreservation and cell culture.

### Generation of single cell suspension for scRNAseq

The distal thrombus tissue isolated from CTEPH patients were cut into ∼ 3 mm pieces and digested using collagenase A for 1 hour. Cells were strained using 70 µM cell strainer. To remove red blood cells, cells were washed with ice cold DPBS and incubated with ACK (red blood cells lysis buffer) on ice for 5 mins. Cells were washed with DPBS and resuspended in cation-free HBSS that contains 10% FBS, 3 mM EDTA and 10 mM HEPES. Viable cells were counted manually using a hemocytometer. Cells were washed in DPBS and stained with Aqua (cell viability dye) (1:500). The stained cells were washed with ice cold DPBS and fixed with 80% of methanol for 15 mins. The viable cells were sorted using flow cytometry. After rehydration of methanol-fixed cells, scRNAseq libraries were prepared using the 10X Genomics Single Cell Protocol in collaboration with the Duke Human Vaccine Institute Genetics Analysis Core. Live cells were barcoded with cell-specific barcodes and unique molecular identifier (UMI) based on the Chromium Single Cell 3’ library with Gel Bead Kit version 2 protocol. ScRNAseq libraries was obtained and RNA sequencing was performed that passed quality control (Novogene).

### Generation of endothelial, smooth muscle cell and myofibroblast primary cells

The approach for isolating cells was similar to that used for cells from PAH lungs.^18^ For isolation of endothelial cells (ECs) from thrombus tissue, the thrombus tissue was cut into 2 cm pieces and then rinsed 3 times with HBSS to remove blood cells. The tissue segments were incubated in 10-15 ml PBS with 2 mg/ml type II collagenase at 37ºC, 5% CO2 with 90% humidity for 20 minutes. Tissue segments were massaged with spatula followed by gentle shaking to detach ECs into endothelial basal media (EBM) media with 1% pen/strep/fungizone. The thrombus tissues were removed and the cells were centrifuged at 330g for 7 minutes at room temperature. Cells were re-suspended in EBM2 media supplemented with growth factors (Lonza). Cells were plated on gelatin-coated cell culture plates. For isolation of smooth muscle cells (SMCs) from thrombus, the resulting tissue segments were digested overnight in 100 ml of HBSS which contains collagenase and DNase (each 0.1 mg/ml), 2.5 ml HEPES buffer, penicillin (250 units/ml), streptomycin (250 mg/ml) and amphotericin B (0.625 mg/ml). After digestion, cells were strained using a 100 um-pore nylon cell strainer. Cells were plated and cultured in SMC growth media. For isolation of myofibroblasts, the tissues were cut into ∼1 mm pieces. The tissues were incubated in 10 mL of DMEM/F12 media with 0.14 Wunsch units/mL Liberase Blendzyme 3, and 1X antibiotic/antimycotic at 37°C for 30 to 90 minutes. After digestion, warm DMEM/F12 media with 15% FBS, 1X antibiotic/antimycotic was added to stop the liberase digestion. The cells were centrifuged at 524g for 5 mins and supernatant was removed. This step was repeated three times to remove the traces of Liberase. The cell pellet was resuspended in 10 mL of DMEM/F12 media with 15% FBS, 1X antibiotic/antimycotic and transfered to a 10 cm tissue culture dish. The myofibroblasts were cultured in a tissue culture incubator at 37°C, 5%CO_2_.

### Immunohistochemistry

Immunohistochemistry for distal thrombus tissue from CTEPH patients was performed by the Department of Pathology, Duke University Medical Center. Thrombus tissue sections were stained for Hematoxylin and eosin stain (HE) and Masson’s trichrome (MT) staining. Images were taken using Zeiss Axio Imager Widefield Fluorescence Microscope.

### Immunofluorescence

Cells isolated from thrombus and control human pulmonary artrery were grown on chamber slides. Cells were washed and fixed with 4% PFA for 15 mins at room temperature. Cells were permeabilized with 0.2% Triton X-100 for 5 mins at room temperature. Cells were blocked with 5%BSA for 1 hour. Primary antibodies were incubated at 4°C for overnight. Cells were washed and incubated with DAPI for few mins. Images were acquired with Leica SP5 Inverted Confocal using Leica LAS AF 2.6 software at 40X magnification. Antibodies used in this study were: vWF-Alexa Fluor 488 (Abcam Cat. 195028), αSMC actin-Cy3 (Sigma Cat. C6198), vimentin-Alexa Fluor 647 (BioLegend Cat. 677807), FSP1-FITC (BioLegend Cat. 370007). Tissue slides were made from OCT blocks of distal thrombus tissue from CTEPH patients. Samples were stained with cell specific markers for macrophages (CD206), T-cells (CD3), endothelial cells (vWF), smooth muscle cells and myofibroblasts (αSMC actin). Tissue slides were mounted with DAPI mounting media (Invitrogen Cat. P36931). Images were acquired with Leica SP5 Inverted Confocal using Leica LAS AF 2.6 software at 40X magnification. Antibodies used in this study were: vWF (Invitrogen Cat. MA5-14029), CD206 (BioLegend Cat. 141701), CD3 (Invitrogen Cat. MA1-21454) amplified with TSA Cy3 kit (PerkinElmer Cat. FP1170), αSMC actin-Alexa Fluor 660 (Invitrogen Cat. 50-9760-82).

### Tube formation assay

The ability of CTEPH ECs to form tube-like structures on basement membrane matrix was compared to control human pulmonary artery endothelial cells (PAECs) obtained commercially (Lonza). The Cultrex® Reduced Growth Factor Basement Membrane Matrix (Trevigen cat.no. 3433-001-01) was added to wells of a 96 well plate (30μl/well). The matrix was allowed to polymerize for 30 mins at 37°C and 5% CO_2_. Cells were plated onto the basement membrane matrix coated plates. After 24 hours, the cells were imaged under an inverted microscope at low magnification and number of tubes formed were counted manually.

### Proliferation assay

Cell proliferation was assessed using a Cell Proliferation ELISA, BrdU (colorimetric) Kit (Roche Applied Science, Indianapolis, IN). Cells were plated on flat bottom 96 well plates (5000 cells per well) in complete growth medium. The cells were labeled using 10 μM BrdU per well and re-reincubated for 6 hours at 37°C in incubator. The culture medium was removed, cells are fixed, and the DNA was denatured using FixDenat for 30 minutes. The cells were incubated with the anti-BrdU-POD antibody for 90 minutes at room temperature. After the removal of the antibody, the cells were washed and the substrate was added. The chemiluminescence was quantified by measuring the absorbance at 370nm and 492nm using a BioTek Synergy Neo 2 multi-mode reader. For studies with PAR1 inhibition, cells were starved for 6 hours, stimulated with or without 10 μM α-thrombin and/or the PAR1 antagonist vorapaxar 10 μM for 24 hours.

### Apoptosis assay

Apoptosis was assessed using a terminal deoxynucleotidyl transferase dUTP nick-end labeling (TUNEL) assay (HT TiterTACS™ assay kit, Trevigen, Gaithersburg, MD) according to manufacturer’s instructions. Briefly, 2 × 10^4^ CTEPH and control cells were cultured on a 96 well plate for 24 hours. The cells were washed with PBS and fixed with 3.7% formaldehyde solution for 7 mins at room temperature. Cells were washed with PBS and incubated in ice cold methanol for 20 mins. To quench endogenous peroxidase activity, cells were treated with 3% hydrogen peroxide solution for 5 min and washed with distilled water. The cells were incubated in labeling reaction mix for 1 hour at 37°C, and the reaction was stopped by incubation in 1x TdT stop buffer for 5 minutes. The cells were washed with PBS and incubated with Strep-HRP for 10 mins. After that incubation, the TACS Sapphire solution was added for 30 minutes at room temperature. TUNEL positive cells were measured at 450 nm of absorbance using a BioTek Synergy Neo 2 multi-mode reader. As a positive control, cells were treated with TACS-nuclease provided in the assay kit for 1 hour at 37°C before hydrogen peroxide treatment.

### In vitro scratch assay

Cells were plated on 24 well plates. At 80% confluence, cells were starved for 6 hours and scratches were made using a 20 μl pipette tip. The cells were washed and stimulated with or without 100 nM α-Thrombin and 1 μM vorapaxar. Wound closure was monitored using a live-cell station Zeiss Axio Observer microscope (Duke Light Microscopy Core Facility). The images were captured in real-time at 0 hour and for every hour for 12 hours. The initial edges of the scratch at 0th hour was marked and migrated distance at 12 hours was measured using MetaMorph Premier (Molecular Devices).

### Statistics

All graphs and data generated in this study were analyzed using GraphPad Prism 9 Software. All quantitative data is presented as means ± SEM. The statistical significance of differences was determined using Student’s two-tailed t test in two groups, and one-way or two-way ANOVA along with Bonferroni, Tukey’s or Sidak’s multiple comparison test in multiple groups. Survival curves were analyzed by the Kaplan-Meier method and compared by a log-rank test. A p value < 0.05 was considered statistically significant.

### Single-cell RNA-seq analysis

#### Mapping of reads to transcripts and cells

The sequencing data was processed into transcript count tables with the Cell Ranger Single Cell Software Suite v3.0.1 (10X Genomics). The FASTQ files for each of the five CTEPH samples were processed independently with the *cellranger count* pipeline to build a transcript count table for each sample. This pipeline performs the following steps: 1) STAR^19^ is used to align cDNA reads to the Homo sapiens transcriptome (Sequence: GRCh38, Annotation: Gencode v93); 2) cell barcodes and unique molecular identifiers (UMIs) corresponding to aligned reads are then filtered and corrected; and 3) reads associated with retained barcodes are quantified.

#### Data integration

The resulting transcript count tables were analyzed using the R *Seurat* package v4.0.0^20^. Standard preprocessing (log-normalization) was performed individually for each of the five CTEPH samples. Variable features (nfeatures = 2000) were identified based on a variance stabilizing transformation (selection.method = “vst”). We then identified anchors using the FindIntegrationAnchors function, which were subsequently used for integration with the method implemented in the IntegrateData() function with default parameters.. The new integrated matrix was then used for downstream analysis and visualization.

#### Joint clustering and cell type annotation

To exclude low quality cells, we filtered out cells for which fewer than 300 UMIs or 200 genes were detected. Likely doublets were excluded by removing cells with greater than 60,000 UMIs or 9,000 genes detected. We also filtered out cells for which the percent of mitochondrial gene expression represented more than 15% of the total gene expression. The dataset ultimately consisted of 11,709 cells. Sample CTEPH-B and CTEPH-E were of lower quality both in terms of number of cells (132 and 225 respectively) and QC metrics such as the median number of genes detected per cell (590 and 336 respectively, compared to >967 cells for the three other samples). We then used principle component analysis and identified 30 principle components (PCs) for downstream analysis. The shared nearest neighbor (SNN) modularity optimization-based clustering algorithm implementec in the FindClusters function in *Seurat* were used to identify cell clusters with a resolution of 0.6. In general, resolutions for clustering were chosen aided by the clustree method^21^ to visualize how clusters breakdown and how selected marker genes are expressed in these clusters from one resolution to another. Fifteen distinct clusters of cells were identifified and visualized in two dimensions using nonlinear dimensionality-reduction technique with uniform manifold approximation and projection (UMAP). The *Seurat* FindConservedMarkers function (Wilcoxon rank sum test) was used to identify cluster-defining differential expressed genes (DEGs). The R package *SingleR* v1.4.0^22^ was used to annotate cell types based on correlation profiles with a collection of Affymetrix microarrays deposited in the Gene Expression Omnibus (GEO) repository, and combined into the Human Primary Cells Atlas (HPCA)^23^.

#### Macrophages and T-cells subclustering

Based on previously identified clusters and after removing lower quality samples CTEPH-B and E, we took subsets of the data set corresponding to macrophages (n = 1,846 cells),, T-cells (n = 3,037 cells cells), and smooth muscle cells (SMCs, n = 4,751 cells). For each of these subsets, we followed a similar workflow as for the whole data set to integrate and then cluster. For macrophages, we performed a first clustering (resolution 0.2) that identifies five clusters. One of these clusters (n = 109 cells) was distinct from the others and identified as residual T cells. We thus performed a second clustering after filtering out this cluster and selected a resolution of 0.3 which identified five clusters. Similarly, for the T-cells subset, we performed a first clustering (resolution 0.2) that identified five clusters. One of these clusters (n = 110 cells) was identified as mesenchymal cells. We then performed a second clustering excluding this cluster that identified four clusters (resolution 0.2). For the mesenchymal cell subset, resolutions were chosen based on the expression levels of COL3A1 as a defining marker for cell populations of interest. The mesenchymal cell subset was first clustered with a resolution of 0.8 that identified nine clusters. One of these clusters (n = 457 cells) appeared to be a technical artefact clustering on high mitochondrial content. Another one was identified as immune cells. We then performed a second clustering (resolution 0.7) after removing these two clusters which identified eight clusters.

#### Smooth Muscle Cell subclustering and Pseudotime Analysis

To infer the lineage relationships between SMC sub-clusters, we used approaches implemented in the R package *Monocle3*^*24*^ (https://github.com/cole-trapnell-lab/monocle3). Based on the expression of marker COL3A1, one of the subclusters was identified (and labeled) as ‘Myofibroblasts’. Other subclusters were labeled ‘SMC1’ to ‘SMC7’. After re-normalization of the raw count data by log and size factor and dimensional reduction by PCA, sample SMC subsets were aligned using a mutual nearest neighbor method named Batchelor described in Haghverdi, et al. ^25^ and implemented in *Monocle3*. The UMAP was then calculated for the SMC subset and showed that the smallest cluster SMC7 (n = 71 cells) contained outlier cells that would not be included in the main trajectory. We therefore decided to remove SMC7 to focus on the main trajectory and redid the initial steps. *Monocle3* partitions the cells into supergroups called partitions using a method derived from “approximate graph abstraction” (AGA) described in Wolf, et al. ^26^. Cells from different partitions cannot be part of the same trajectory. Only one partition was detected in the 4,102 remaining SMCs. Trajectory graph learning and pseudo-time measurement were performed with *Monocle3*. Principal graph was learned from the reduced dimension space using reversed graph embedding and setting the number of graph nodes at 200.

Single-cell trajectory was rooted using cells from subset ‘SMC2’ based on high expression of the ACTA2 marker in that cluster, which allowed the ordering and generation of pseudotime values for every cell in the data set. To identify genes that change as a function of pseudotime, we used an approach based on the Moran’s I test as implemented in *Monocle3* (q-value < 0.05). To uncover potential gene co-regulation networks, we then used the *find_gene_modules* function of *Monocle3* with default parameters. This approach clusters genes into modules that are co-expressed across cells. We selected six modules of interest both based on SMCs markers and the max and min levels of aggregated gene expression in the “Myofibroblasts” cluster. The genes belonging to each module were used to run Core Analyses in Ingenuity Pathway Analysis to help characterize each module. The lineage leading from cluster ‘SMC2’ to ‘Myofibroblasts’ was selected by using the *choose_graph_segments* function. The gene expression along pseudotime data of several markers and genes of interested were then visualized using the *plot_genes_in_pseudotime* function.

#### Pathway Analyses

To characterize subclusters from T cells, macrophages and mesenchymal cells, as well as gene modules identified by the single-cell trajectory analysis of mesenchymal cells in the context of biological pathways, the ‘Core Analysis’ included in the Ingenuity Pathway Analysis software (IPA, Ingenuity System Inc, USA, http://www.ingenuity.com) was run on differentially expressed genes (DEGs) for each analysis. For subclusters, the FindAllMarkers function from Seurat was run on all genes and DEGs were selected using an adjusted p-value < 0.05 and and absolute value of log fold-change > 0.25. We used the activation z-score calculated by the ‘Canonical Pathways’ downstream analysis. It predicts whether the upstream regulator exists in an activated or inactivated state. It is a statistical measure of correlation between relationship direction and gene expression. z-score > 2 or < -2 is considered significant. The z-score cannot be calculated for all pathways or gene sets if there is insufficient evidence in the IPA Knowledge Base for confident activity predictions across datasets or if there are fewer than four analysis-ready molecules in the dataset associated with the pathway.

#### Cell type Ligand-Receptor Interaction Analysis

Ligand-receptor interactions were calculated between one set of ligand-expressing cells (*l*) and another set of receptor-expressing cells (*r*). The ligand-receptor interaction score was calculated as the average product of ligand and receptor expression across all single-cell pairs (*l*_*i*_, *r*_*j*_) between the ligand-expressing cell set and the receptor-expressing cell set

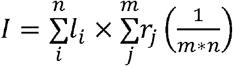

Where *I* is the ligand-receptor interaction score, *n* is the total number of cells in the ligand-expressing cell set, and *m* is the total number of cells in the receptor-expressing cell set.

Prior knowledge of the ligand and receptor interactions was obtained from Ramilowski et al.^27^. The cell types that were chosen for the analysis were the T cells, macrophages, and smooth muscle cell (including myofibroblast) clusters prior to any subclustering. Potential ligand-receptor interactions were calculated for these three cell types, resulting in nine cell-type/cell-type interaction scores. Visualization of the interactions was performed using CIRCOS plots (http://mkweb.bcgsc.ca/tableviewer/).

## Results

### Study Design and Methods

CTEPH thrombus samples had a significant cellular component, which we confirmed with histopathology of distal thrombus samples (**Fig. 1A** and **B**). These samples demonstrated significant fibrosis with areas of recanalization with a significant cellular component. These samples also demonstrated the presence of CD206+ macrophages (**Fig. 1C**), CD3+ T cells (**Fig. 1D**), consistent with the prior observation of immune cell infiltration of CTEPH thrombus (**Supp. Fig. 3**). Populations of ECs that expressed vWF, and SMCs that expressed a-SM actin were also observed(**Fig. 1E**), ^13^ Based on this, we processed five distal CTEPH thrombus samples for scRNAseq analysis (see *Materials and Methods*). **Supplementary Table 1** summarizes the demographic and clinical characteristics of the patients whose thrombus removed during PTE surgeries was used for these studies. Preoperative imaging (computed tomography angiography of the chest and ventilation-perfusion scan) for three of those five subjects is shown in **Supp. Fig. 1**. Thrombus removed at the time of surgery for those five subjects is shown in **Supp. Fig. 2**.

**Figure 1.**
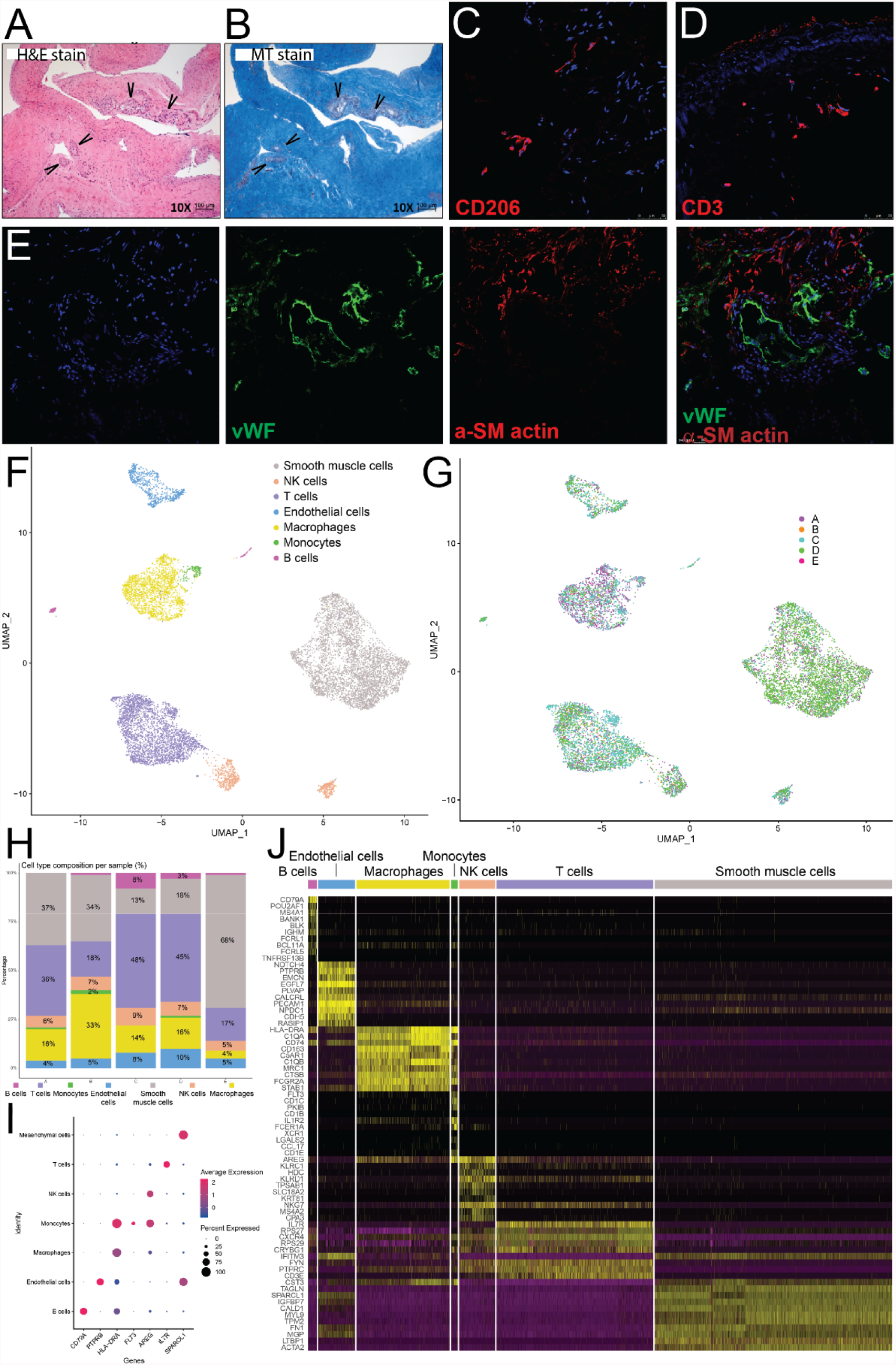
A Cell Atlas of CTEPH thrombus. (**A**) Hemotoxylin and eosin and (**B**) Masson’s trichrome staining of distal PTE samples demonstrate significant fibrosis, recanalization (arrows) and cellularity. Immunofluorescence staining with DAPI (blue) and (**C**) CD206 (red) for macrophages, (**D**) CD3 (red) for T cells, and (**E**) vWF (red) and α-smooth muscle cell actin (purple) for endothelial cells and smooth muscle cells, respectively. (**F**) UMAP plot depicts transcriptome of cells isolated from 5 distal CTEPH thrombus samples. Each dot represents a single cell (n = 11,709). Coloring is according to distinct populations of SMCs, ECs and immune cells identified by the unsupervised clustering performed with *Seurat*. (**G**) These different cell types were found across all five patients (A-E), (**H**) but with distinct percentages of cell types identified in each sample. Some samples were SMC-predominant, while others were T cell- or SMC-predominant. (**I**) Top markers for each cell type. (**J**) Heat map of top ten markers for each cell cluster.

### A Cell Atlas of CTEPH Thrombus

ScRNAseq of CTEPH thrombus identified distinct populations of cells based on annotations from the Human Primary Cell Atlas on the UMAP plot. The largest populations of cells were SMCs and immune cells. There were also minor populations of ECs. (**Fig. 1F**). Across all samples, of the 12,354 cells sequenced from distal regions of CTEPH thrombi, 6,479 cells (52.1%) were immune cells, consisted mainly of T cells (50.7%) and macrophages (32.9%) (**Fig. 1F**). Minor populations of NK cells (11.4%), B cells (2.9%), and monocytes (2.0%) were also observed. These different cell types were present across all patients (**Fig. 1G**), although with different proportions across patients (**Fig. 1H**). Samples from two of the five patients (C and D) had low proportions of SMCs (13% and 18%) but high proportions of T cells (48% and 45%, respectively). Two samples (A and B) had a relatively even distribution of SMCs but differing contributions of macrophages and T cells. One sample (E) had a very high proportion of SMCs (68%), few T cells (17%), and even fewer macrophages and other immune cells. Endothelial cells constituted 10% or less of all cells in all samples. The top markers for each cell type (**Fig. 1I**) and the top ten markers for each cell type (**Fig. 1J**) demonstrated excellent segregation between different cell types. Together, these findings demonstrate that immune cells are a significant component of CTEPH thrombus, consistent with inflammation playing a central role in its pathobiology.

### Macrophages with an inflammatory and remodeling profile are the predominant myeloid cells within chronic thrombus

We then sought to further characterize the major cell types that comprise CTEPH thrombus. Analyzing cells identified as macrophages and applying unsupervised *Seurat*-based clustering, seven macrophage subclusters (Clusters 1-7) were identified (**Fig. 2A**). The separation of these macrophage subclusters was illustrated well by the differential expression of their top markers (**Fig. 2B**) and their cluster-defining differentially expressed genes (DEG) were identified (**Fig. 2C**). Notably, the separation of these clusters did not correlate well with canonical markers of macrophage identity (**Supp. Fig. 4**). Hierarchical clustering of signaling pathways identified by Ingenuity Pathways Analysis (IPA) demonstrated distinct patterns among the DEGs in these clusters (**Fig. 2D**). Cluster 1, 2, and 6 consisted of macrophages enriched in pathways for both classically activated (i.e. HMGB1, INOS, production of NO and reactive oxygen species) with pro-inflammatory features and alternatively activated (STAT3, TREM1 pathways) with tissue remodeling potentials (i.e. cardiac hypertrophy, epithelial to mesenchymal transition, and hepatic fibrosis) (**Fig. 2D**). These three clusters have overlapping cytokine signaling profiles. Cluster 1 and 2 exhibit strongest activation of IL-17, IL-1, IL-6, and IL-8 signaling pathways. Cluster 6 showed predominant activation of IL-8 signaling. In addition to inflammatory cytokine signaling, Fc receptor-mediated phagocytosis pathway was upregulated in cluster 2; and cluster 6 is strongly enriched for autophagy pathway. Thus, cluster 1, 2, and 6 macrophages have pro-inflammatory and pro-remodeling profiles that can establish or perpetuate a chronic inflammatory microenvironment and modulate vascular and thrombus remodeling. On the other hand, clusters 3, 4, and 5 have anti-inflammatory profiles that were enriched in PPAR signaling, and down-regulation of proinflammatory cytokine pathways. Cluster 7 represent a cluster of rare macrophages undergoing cellular division. Thus, there were heterogeneous population of macrophages in chronic thrombus with overlapping functional profiles. The majority of macrophages exhibited pro-inflammatory and vascular remodeling signatures, with minor populations with anti-inflammatory properties.

**Figure 2.**
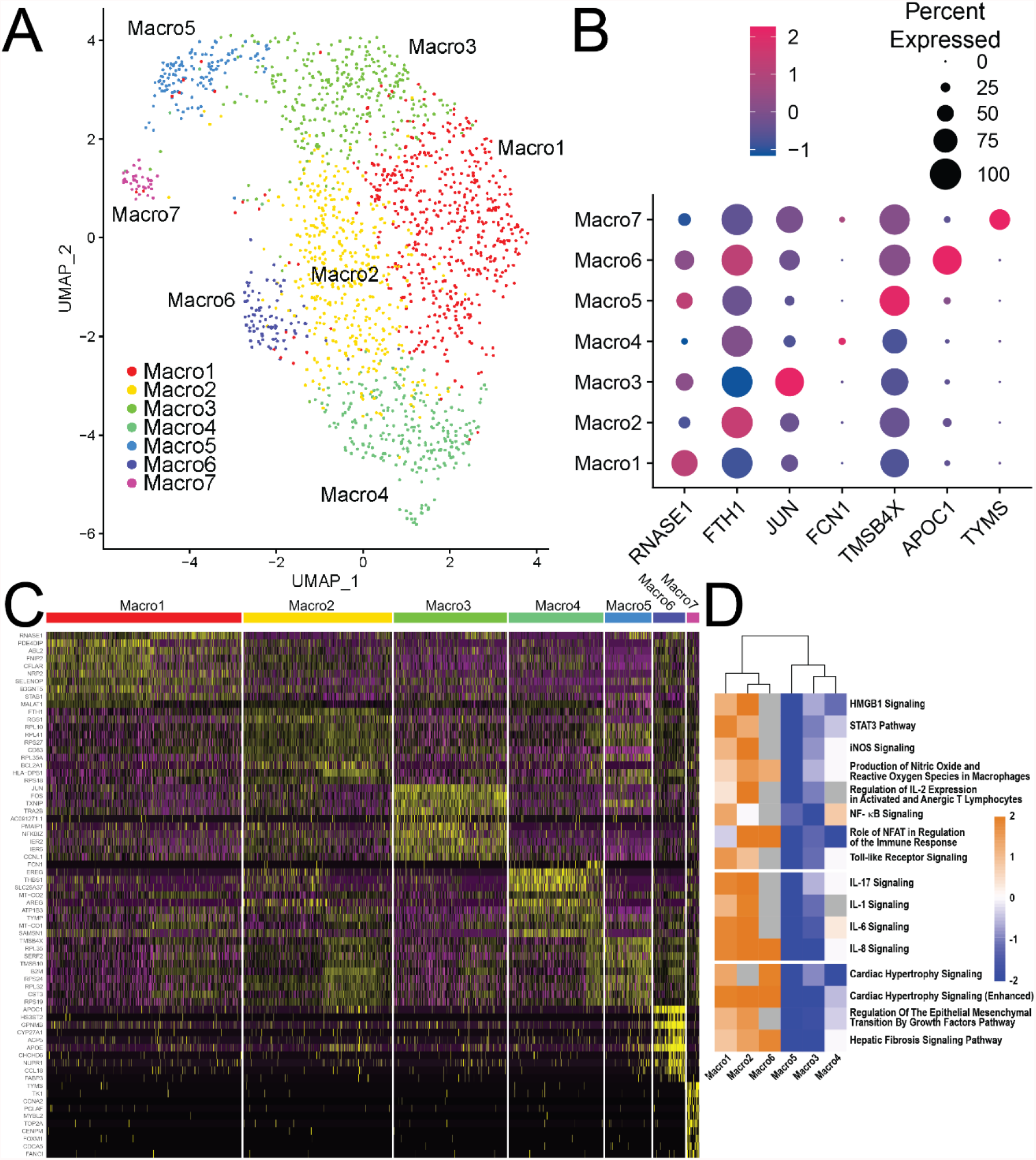
Macrophage clusters in CTEPH. (**A**) UMAP plot of macrophages with their clustering into 7 distinct groups. (**B**) Top markers for each subcluster. (**C**) Heatmap of top ten markers for each subcluster. (**D**) Hierarchical clustering of IPA canonical pathway analysis for macrophage clusters identifies two larger populations of macrophages based on their signaling profiles, with clusters 1, 2 and 6 displaying a pro-inflammatory phenotype.

### Heterogeneous populations of T cells that promote chronic inflammation and autoimmunity are found within chronic thrombus

Analyzing cells identified as T cells and applying Seurat-based clustering analyses, four clusters were identified and annotated based on their expression profiles (**Fig. 3A**). Using *SingleR* in Seurat along with correlation of cell cluster profiles to the HPCA and knowledge-based analysis of DEGs (**Fig. 3B** and **C**), the T cells were clustered as CD4+ T cells, CD8+ T cells (1 and 2), and dividing T cells. IPA canonical pathways analysis identified distinct processes regulated by these clusters (**Fig. 3D**). The CD4+ T cells expressed *CCR7, IL7R, FOXO1, FOXP1*, and *FOXP3*, consistent with effector/memory T regulatory (Treg) cells. This cell cluster also upregulated CTLA-4 and STAT3. STAT3 is known to induce CD4+ Treg cells to produce IL-17,^28^ thereby promoting chronic inflammation. Consistent with this, IPA analyses showed an upregulation of the Th17 activation pathway. Thus, the CD4+ T cell within chronic thrombi are likely Th17-biased Treg cells that uregulate CTLA-4 expression, which have been shown to promote autoimmune diseases and cancer. CD8+ T cell cluster 1 expressed granzyme B (*GZMB*) and CD7, consistent with memory CD8+ cytotoxic T lymphocytes (CTLs). CD8+ T cell cluster 2 expressed granzyme K (*GZMK*) and *CXCR3*, with a gene expression profile consistent with a population of short-lived, granzyme K+ effector CD8+ T cells. Granzyme K CD8+ T cells have been shown to have a limited role in cytotoxicity with a larger role in promoting inflammation through cytokine production (*e*.*g*., *CCL3, CCL4, CCL5, IFNG*, and *TNF*) that can further recruit other immune cell to the site of inflammation.^29^ Additionally, CD8+ T cells cluster 2 also expressed *IL23A*, which can induce IL-17 production by Foxp3+ Treg cells.^30^ Thus, chronic thombi contain a diverse T cell populations that can promote chronic inflammation and autoimmunity.

**Figure 3.**
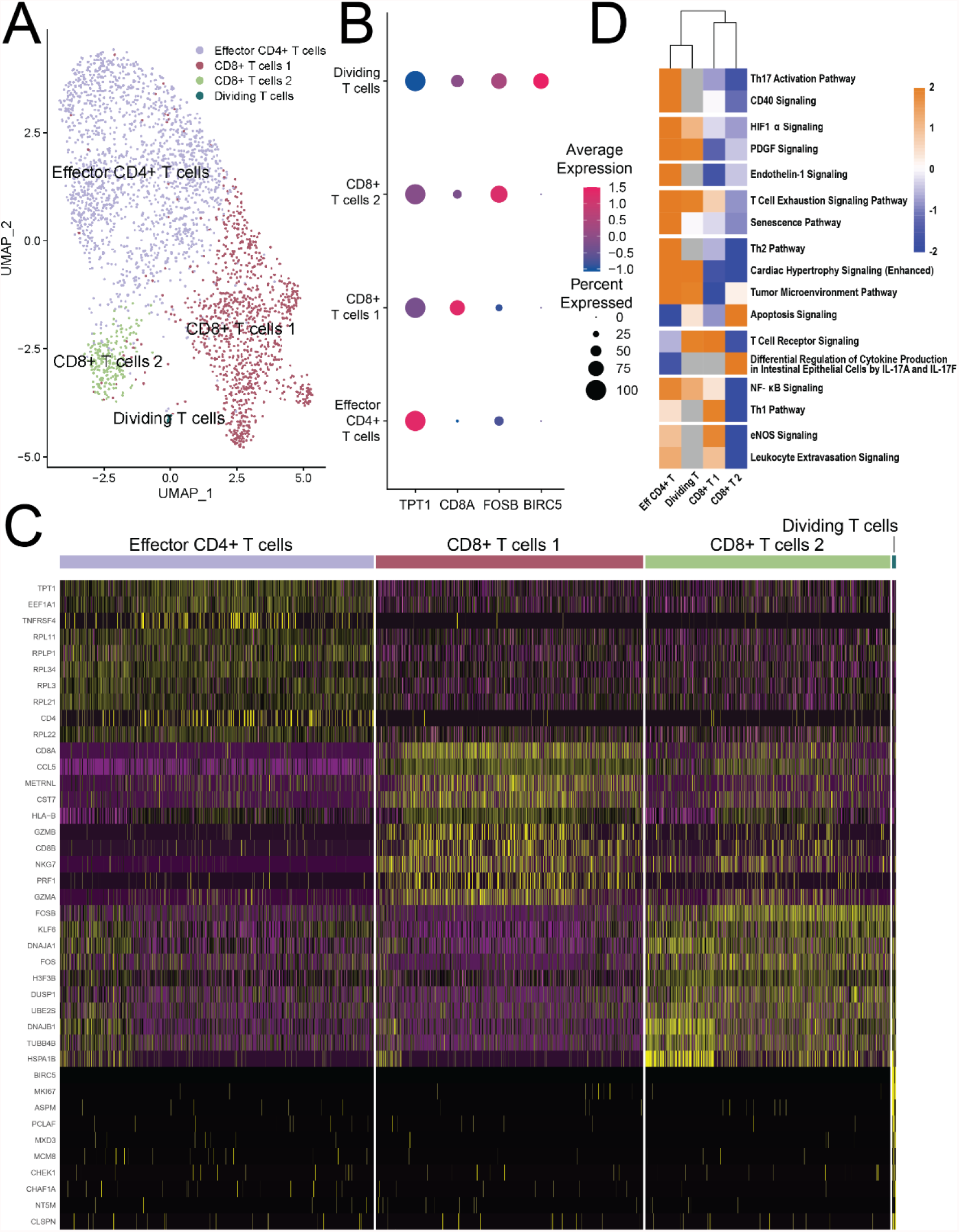
T cell clusters in CTEPH. (**A**) UMAP plot of T cells with their clustering into 4 distinct groups of effector CD4+ T cells, two clusters of CD8+ T cells, and a small cluster of dividing T cells. (**B**) Top markers for each subcluster. (**C**) Heatmap of top ten markers for each subcluster. (**D**) Hierarchical clustering of IPA canonical pathway analysis for T cell clusters.

### Multiple clusters of smooth muscle cells are present in CTEPH thrombus

Analyzing cells identified as SMCs and applying *Seurat*-based clustering analyses, eight clusters of cells wre identified and annotated based on their expression profiles (**Fig. 4A**). One of these clusters was identified as a population of myofibroblasts based on the expression of *COL3A1, COL1A2, DCN* and *LUM* (**Fig. 4B-C**). A small subcluster (SMC7) shared expression of myofibroblast markers but also expressed markers of stem and hematopoietic cells (*CD34*) and consisted only of 71 cells (compared to 515 myofibroblast cells), so it was not studied further. We identified the SMC2 population as a contractile SMC phenotype, as it had the highest expression of *ACTA2* (α-smooth muscle cell actin). IPA analysis of SMC2 also identified the upregulation of signaling pathways important for contractile function including actin cytoskeleton, integrin, Rac and RhoA signaling (**Fig. 4D**). SMC1 also expressed *ACTA2, MYH11* and other markers seen in contractile SMCs, but also ribosomal proteins (*RPL23A, RPL37A, RPL19*, etc.), consistent with SMCs with a proliferative phenotype. While adjacent to SMC2 on the UMAP plot, SMC1 cells displayed a distinct profile on IPA analysis (**Fig. 4D**). Based on their IPA profiles, there was significant heterogeneity across the entire SMC cluster.

**Figure 4.**
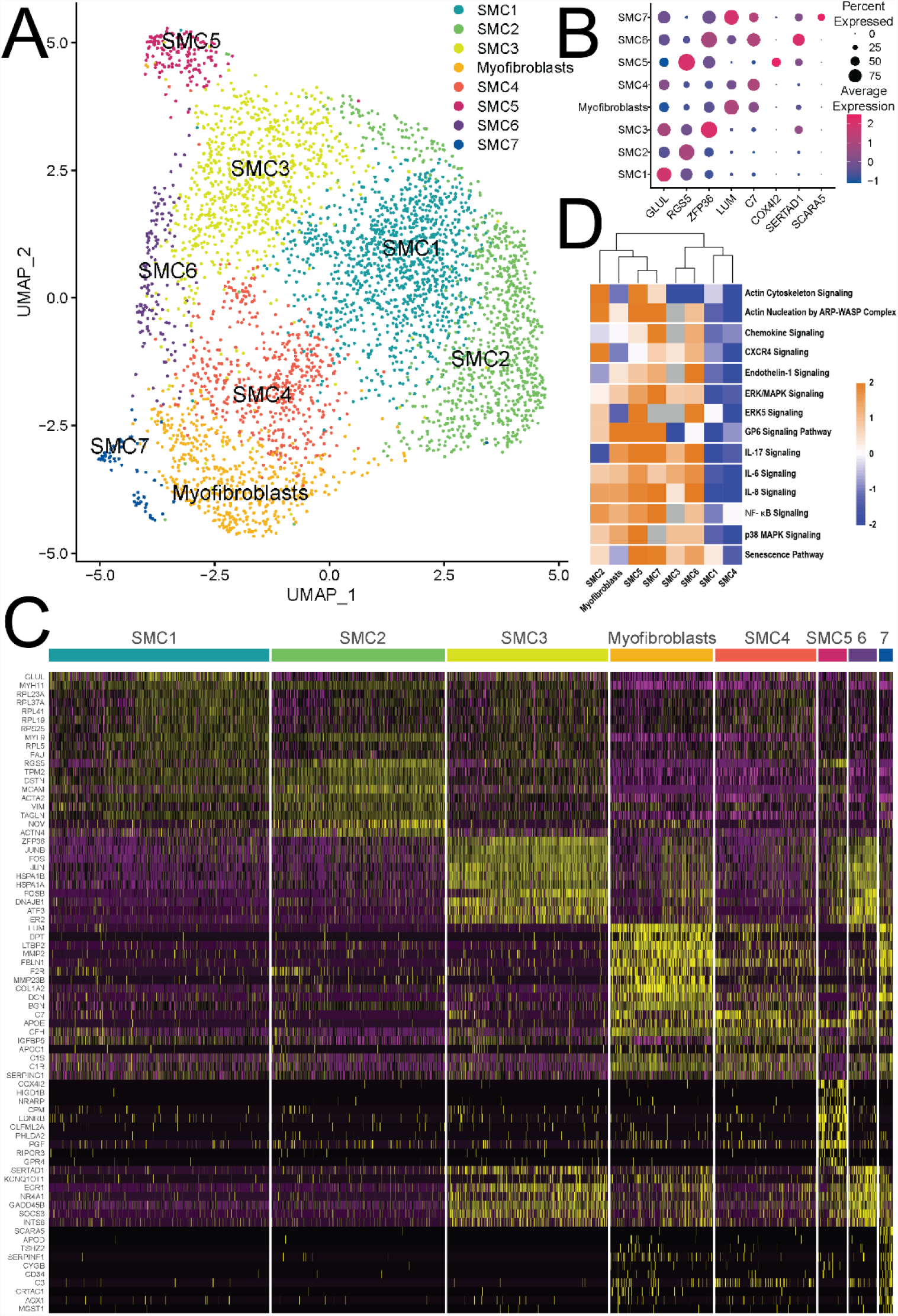
Smooth muscle cell clusters in CTEPH. (**A**) UMAP plot of mesenchymal cells with their clustering into 8 distinct groups. (**B**) Top markers for each subcluster. (**C**) Heatmap of top ten markers for each subcluster. (**D**) Hierarchical clustering of IPA canonical pathway analysis for SMC clusters demonstrates significant heterogeneity in signaling between SMC subclusters.

### Myofibroblasts and SMCs with an inflammatory phenotype likely arise from contractile and proliferative SMC clusters

With the patterns of inflammation and fibrosis seen in some SMC subclusters, we then used pseudotime analysis to test whether myofibroblasts could arise from the SMC1/2 clusters. We focused on the contractile SMC2 cluster as the source of SMCs in CTEPH thrombus. Pseudotime analysis based on *ACTA2* expression identified trajectories from SMC2 to SMC1 and from SMC2 to myofibroblasts (**Fig. 5A** and **B**). These two trajectories likely represent the transition from a contractile to a proliferative phenotype and from contractile SMCs to myofibroblasts. The markers whose expression changed the most on those trajectories were *MYH11, LUM, TPM2* and *MYL9* (**Fig. 5C**). Notably, *MYH11, TPM2* and *MYL9* are all contractile proteins, consistent with a loss of the contractile phenotype over both trajectories. *LUM* is a proteoglycan highly expressed by myofibroblasts^31^. Focusing on the SMC2-Myofibroblast trajectory, we found that the pseudotime analysis predicted a gradual transition from the the contractile SMC2 cluster through clusters SMC1, 3 and 4 until it reached the myofibroblast cluster (**Fig. 5D**). An analysis of gene modules (**Supplementary Table 2**) that were differentially expressed across the pseudotime analysis by SMC cluster also provided important insights from a knowledge-informed analysis (**Fig. 5E**). SMC2 demonstrated high expression of modules 4 (cytoskeletal signaling) and 30 (cytoskeleton/integrins). SMC3 and 6 had high expression of module 6, consisting of genes such as *IL6* and *CCL2* that are important in inflammation. The myofibroblasts had relatively high expression of modules 5, 8, 16, which correspond to genes important in fibrosis (e.g., *TGFB1*), coagulation, complement system, and migration guidance, respectively. Lastly, we analyzed markers of SMC and myofibroblasts to look at their specific expression across the pseudotime trajectory (**Fig. 5F**). As expected, *ACTA2* expression decreased across the trajectory, while myofibroblast markers *LUM, POSTN*, and *COL3A1* all increased significantly. These findings are broadly consistent with SMC clusters with inflammation (SMC3/6) and fibrosis (myofibroblasts) arising from contractile (SMC2) and proliferative (SMC1) SMC clusters.

**Figure 5.**
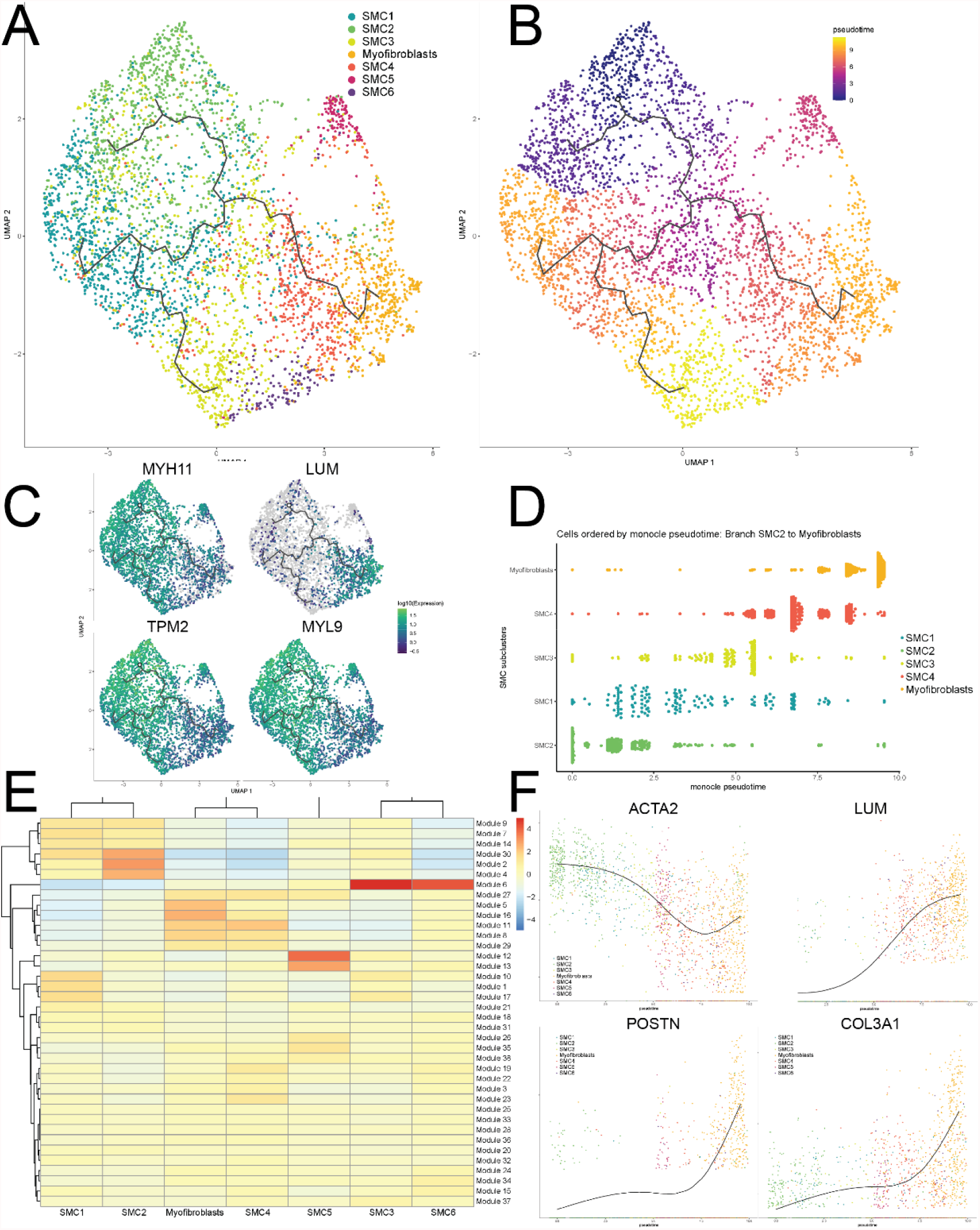
Pseudotime analysis of smooth muscle cell transition to myofibroblasts. (**A**) Pseudotime trajectory from the *ACTA2*-high expression SMC2 subcluster to myofibroblasts and the SMC1 subcluster. (**B**) Cells colored by pseudotime, demonstrating that myofibroblasts and the SMC3 subcluster, which expressed inflammatory genes, arise late in the pseudotime trajectory. (**C**) Four markers that demonstrated some of the largest changes over the pseudotime trajectory, consistent with a loss of contractile protein expression (*MYH11, TPM2* and *MYL9*) and a gain of myofibroblast markers (*LUM*). (**D**) Cells ordered by monocle pseudotime across the SMC2 to myofibroblast trajectory. (**E**) Specific gene modules that were enriched in each subcluster (for details on genes in each module, see **Supplementary Table 2**). (**F**) Loss of expression of *ACTA2* along the trajectory from SMC2 to myofibroblasts, along with an increased expression of myofibroblast markers *LUM, POSTN* and *COL3A1*.

### CTEPH thrombus-derived cells have a distinct phenotype from control pulmonary artery cells

EC, SMC and myofibroblasts were isolated and cultured from CTEPH thrombus as described in *Materials and Methods*. These cells were first compared to cells isolated from control donors (pulmonary artery (PA) ECs (PAECs), PASMCs, and PA adventitial fibroblasts (PAAFs)). Both CTEPH ECs and PAECs expressed vWF in the absence of SMC or mesenchymal cell markers. CTEPH SMCs and PASMCs expressed both α-SM actin and vimentin, but did not express the EC-specific markers vWF (**Fig. 6A** and **Supp. Fig. 5**). Similarly, both CTEPH myofibroblasts and PAAFs expressed both α-SM actin and vimentin in the absence of vWF expression, and did not differ significantly in appearance from SMCs. These findings are largely consistent with our scRNAseq data, which demonstrated significant overlap between the SMC/myofibroblast phenotype.

**Figure 6.**
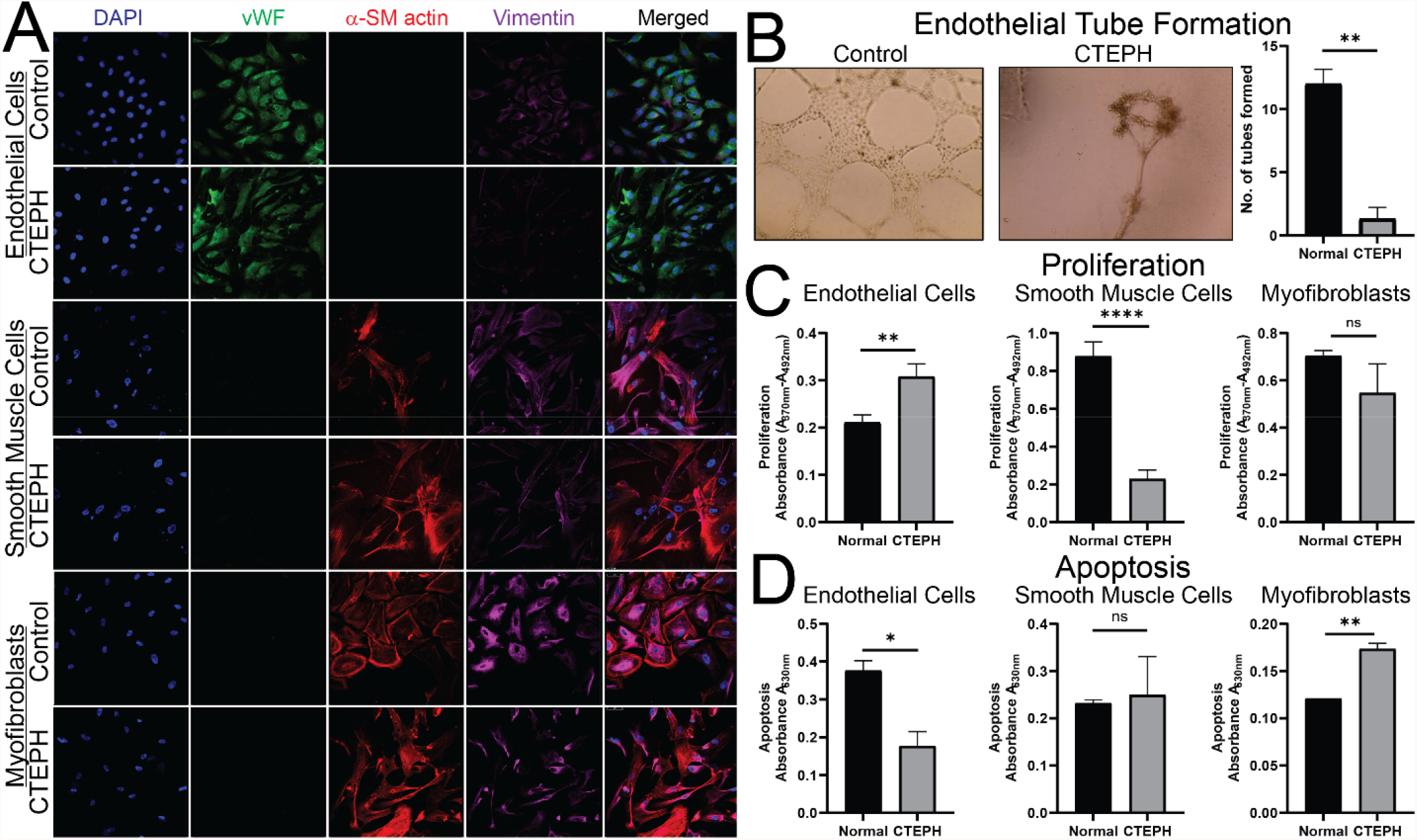
CTEPH thrombus-derived cells have distinct phenotypes from control cells. (**A**) Immunofluorescence with markers for endothelial cells (vWF), smooth muscle cells (α-SMC actin), and mesenchymal cells (vimentin). Compared to PAECs, CTEPH ECs demonstrated (**B**) abnormal tube formation, along with (**C**) increased proliferation and (**D**) decreased apoptosis. Compared to PASMCs, CTEPH SMCs displayed (**E**) decreased proliferation with (**F**) similar levels of apoptosis. Compared to PA adventitial fibroblasts, CTEPH myofibroblasts displayed (**G**) similar levels of proliferation with (**H**) increased levels of apoptosis.

We then assessed the functional properties of these cells. We first tested the ability of CTEPH ECs and PAECs to form tubes. While PAECs were able to form networks of tubes, tubes formed by CTEPH ECs were morphologically abnormal and severely reduced in number (**Fig. 6B**). Consistent with this abnormal phenotype, CTEPH ECs demonstrated significantly higher proliferation than PAECs (**Fig. 6C**), along with a decrease in apoptosis (**Fig. 6D**). CTEPH SMCs demonstrated a significantly different phenotype, with decreased proliferation relative to PASMCs (**Fig. 6E**) with similar levels of apoptosis (**Fig. 6F**). CTEPH myofibroblasts had similar proliferation to PAAFs (**Fig. 6G**) but with higher levels of apoptosis (**Fig. 6H**). Together, these findings are consistent with CTEPH ECs having a phenotype similar to ECs seen in pulmonary arterial hypertension (PAH) (abnormal tube formation, increased proliferation and decreased apoptosis)^32^. This is in contrast to CTEPH SMCs and myofibroblasts, whicih displayed a phenotype that may be related to a loss of proliferation and the expression of inflammatory and fibrotic genes.

### PAR1 Inhibition Reduces SMC and Myofibroblast Proliferation and Migration

We next sought to identify potential ligand:receptor interactions between different cell types (T cells, macrophages and SMC/myofibroblasts) that contribute to vascular remodeling in CTEPH. We performed an unbiased analysis based on expression of potential ligands and receptors on these different cell types and visualized them using CIRCOS plots (**Supp. Fig. 6**). This analysis demonstrated some broad trends, with the potential for GPCR and TGF-β signaling by all cell types and fibronectin/integrin signaling by SMCs, but with no clearly identified targets. We then used a knowledge-based approach to identify potentially important ligand:receptor interactions. As myofibroblasts may promote abnormal pulmonary vascular remodeling, their selective targeting may be a treatment for CTEPH. Based on DEGs, we searched for myofibroblast markers that could serve as drug targets. One of the top ten markers for myofibroblasts was the G protein-coupled recepor *F2R* (**Fig. 4A**), the factor II receptor, also known as protease-activated receptor 1 (*PAR1*). *PAR1* expression was enriched in myofibroblasts (**Fig. 7A**) compared to other SMC clusters, with a significant increase across the SMC-to-myofibroblast pseudotime trajectory (**Fig. 7B**). We then tested the effects of PAR1 activation with thrombin and PAR1 inhibition with the small-molecule vorapaxar on CTEPH SMC and myofibroblast proliferation and migration. In CTEPH SMCs, thrombin promoted cell proliferation, an effect that was inhibited by vorapaxar (**Fig. 7C**). CTEPH myofibroblasts displayed significantly higher proliferation than SMCs in response to thrombin, which was similarly inhibited by vorapaxar (**Fig. 7D**). This difference in proliferation may be due to increased *PAR1* expression in CTEPH myofibroblasts compared to SMCs. The same pattern was observed with migration, where myofibroblasts had an increased response to thrombin compared to SMCs, with both responses inhibited by vorapaxar (**Fig. 7E-F**).

**Figure 7.**
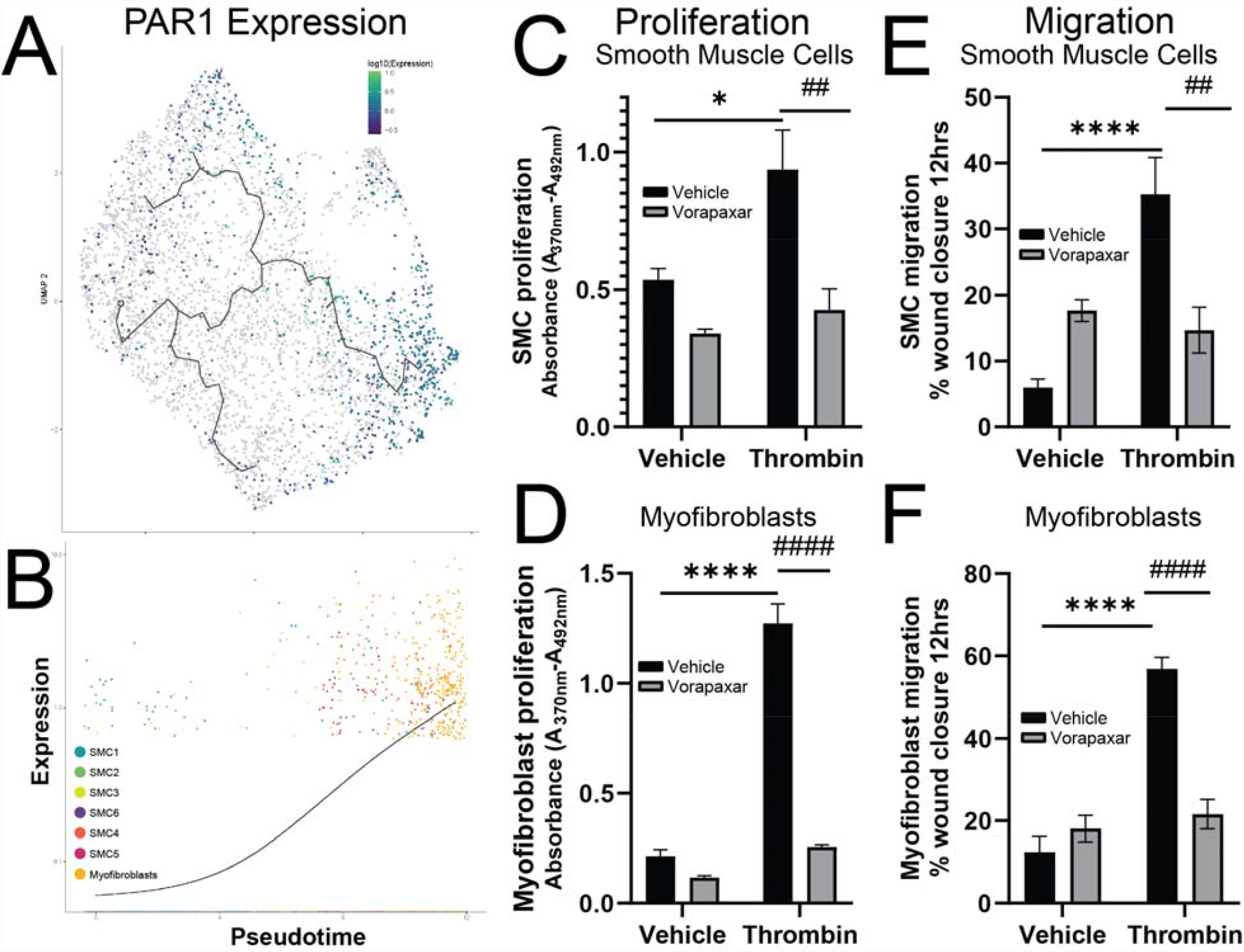
PAR1 antagonism decreases CTEPH SMC and myofibroblast proliferation and migration. (**A**) *PAR1* (*F2R*) expression is enriched in myofibroblasts compared to other SMC subclusters and (**B**) increases in expression across the pseudotime trajectory. Proliferation of CTEPH thrombus-derived (**C**) SMCs and (**D**) myofibroblasts was promoted by 1 µM thrombin (black bars) and inhibited by treatment with 10 µM of the PAR1 antagonist vorapaxar (gray bars). Migration of (**E**) SMCs and (**F**) myofibroblasts was promoted by 100 nM thrombin (black bars) and inhibited by 1 µM vorapaxar (gray bars) (see **Supp. Fig. 7** for representative images).

## Discussion

In this study, we have identified specific cell types that contribute to pathological remodeling, demonstrating important roles for inflammation and SMC modulation in CTEPH. Previous studies on CTEPH thrombus using specific markers have provided important insights into disease pathophysiology, but limited insights into the heterogeneous cell populations that compose CTEPH thrombus and contribute to its pathobiology.^6^ Our work here clearly demonstrates that distinct populations of ECs, SMCs, macrophages, T cells, NK cells and B cells that contribute to CTEPH thrombus. This is largely consistent with a recent single cell study^33^ followed by pseudotime analysis^34^ that demonstrated similar populations of cells (SMCs, myofibroblasts and immune cells). However, there are a number of novel insights that our approach has now allowed, which may be related to differences in data analysis and the use of CTEPH-derived cells to test the importance of specific pathways. Unlike the previous study, we did not have large percentages of unidentified cells (20%), which may be related to the potential inclusion of doublets and cells with high mitochondrial content or to a reliance on cell identification from databases alone.^33^ Rather, we were clearly able to identify pathophysiologically relevant subclusters of cells within all of our larger clusters of macrophages, T cells and SMCs. Our results strongly argues for a process in which macrophage and T cells provide an inflammatory milieu that promote phenotypic modulation of SMCs to myofibroblasts and pulmonary vascular remodeling. The specific overexpression of PAR1 in the myofibroblasts could serve as a mechanistic link between thrombosis and pulmonary vascular remodeling in CTEPH, and inhibiting PAR1 signaling

For our analysis, we chose to focus on the macrophage, T cell and SMC populations as those were the predominant cell types identified. Notably, we observed significant heterogeneity in these cell populations. However, a common finding in the macrophage and T cell subclusters was a pro-inflammatory phenotype. Consistent with this, we observed subclusters of SMCs that expressed inflammatory markers (SMC3 and 6) as well as a population of myofibroblasts. These findings suggest that ongoing inflammation plays an important role in promoting pulmonary vascular remodeling in CTEPH, a mechanism which has also been proposed in PAH.^35^ It is also likely that the other cell populations also play important roles in CTEPH pathogenesis, but an assessment of their roles in CTEPH pathogenesis is limited by a lack of suitable animal models for the disease.^36^

The immune and SMC cell types we observed are similar to those observed in recent single cell studies of atherosclerosis^37-39^. In a mouse model of atherosclerosis with lineage tracing, a population of modulated SMCs with increased expression of *TCF21* arose from tomato-labeled SMCs with the development of atherosclerosis.^38^ These modulated SMCs with expression of fibrotic markers are consistent with the myofibroblast population identified in our studies. Another single cell study identified SMC-derived intermediate cells that were multipotent and could differentiate into macrophage-like and fibrochondrocyte-like cells as well as return toward the SMC phenotype.^40^ While we observed SMC subclusters that are consistent with modulated populations through the expression of inflammatory or fibrotic markers, further studies will be required to determine whether these cells are truly required for pulmonary vascular remodeling in CTEPH. Similarly, populations of pro-inflammatory macrophages and T cells have been identified in human atherosclerosis,^37,39^ similar to the populations that we identified in this study. These parallels suggest that the same pathogenic processes underlie atherosclerosis and CTEPH, with a response to injury in both (fatty streak in atherosclerosis and PE in CTEPH). In this model, pro-inflammatory macrophages and T cells are recruited to the site of injury, secreting factors that promote SMC modulation to myofibroblasts and SMCs with an inflammatory phenotype, that in turn promote pulmonary vascular remodeling. This is also consistent with CTEPH histopathology, in which neointimal, atherosclerotic and recanalized lesions are observed.^13^

Our findings are also consistent with previous studies that have identified ECs and myofibroblast-like cells in CTEPH thrombus^41,42^. For example, the minor SMC7 cluster we observed in our data is consistent with previously described cells that express CD34 (endothelial marker) and α-SMA (SM-cell marker) in endarterectomized tissues from patients with CTEPH^43^. Based on that expression pattern, it was hypothesized that the microenvironment provided by thromboemboli might promote those putative progenitor cells to differentiate and enhance intimal remodeling ^43^. However, in contrast to our findings, CTEPH ECs have previously been shown to be highly proliferative with a high angiogentic potential.^44^ Here we found that while these cells were highly proliferative and apoptosis-resistant, they displayed markedly decreased angiogenic potential as assessed by tube formation^32^. While we found that SMC and myofibroblast proliferation were not significantly different from controls with respect to proliferation and migration, CTEPH SMCs have previously been shown to have a number of differences from control cells, including less or no cell-cell contact inhibition growth, increased sensitivity to hypoxia-induced proliferation, resistance to serum starvation-induced apoptosis, and changes in mitochondrial metabolism^45^. Similarly, myofibroblast-like cells cultured from CTEPH thrombus have been characterized as hyperproliferative, anchorage-independent, invasive and serum-independent^42^. These differences between cultured cells may reflect real differences between cell types or patients, but may also reflect artifactual changes associated with the isolation, propagation and culture of cells between studies.

As in atherosclerosis, approaches that target pathologic remodeling may be beneficial in CTEPH. We found that *PAR1* was selectively expressed in myofibroblasts in the SMC cluster, although it is also known to be expressed in other cell populations. For example, a subpopulation of PAH patients with increased propensity to thrombotic events have previously been noted to have increased platelet *PAR1* expression^46^. Administration of a PAR1 antagonist in rat and mouse PAH models significantly reduced pulmonary vascular resistance, the development of right ventricular hypertrophy, and vascular remodeling, prolonging survival.^47^ Here our functional assays revealed that PAR1 activation with thrombin promoted proliferation and migration of both CTEPH SMC and myofibroblasts, effects that were attenuated by vorapaxar, a PAR1 antagonist. This suggests that PAR1 may be a drug target in CTEPH, where it may serve as a mechanistic node linking thrombosis and pulmonary vascular remodeling.

There are potential limitations with our experimental strategy. As CTEPH is a relatively rare disease, we were limited in the number of samples, which were obtained from a single center. From our approach to sampling PTE thrombus, we cannot differentiate as to whether the relative proportions of SMCs, T cells and macrophages between samples represents a true difference between patients or differences in sampling between thrombi. While we optimized our scRNAseq protocol to isolate the total number of cells from thrombus, it is possible that those conditions resulted in the loss of specific cell populations. Also, as noted above, our studies with CTEPH cells are limited by culture and propagation conditions that can result in phenotypic modulation. Lastly, a target for medical therapies is small vessel disease in CTEPH that affect pulmonary arterioles.^1^ Our scRNAseq studies focused on distal thrombus at the level of segmental and subsegmental pulmonary arteries, and may not reflect the biology of pulmonary arterioles. This limitation is shared by most studies of PAH, where cells isolated for PAH studies are from the larger vessels and still display an abnormal phenotype,^18^ similar to what we observed in our study.

## Author Contributions

GV, SR and YRY designed the study and wrote the manuscript. AG and SAP helped with patient consent. AG and JH performed PTE surgery and thrombus isolation. SR reviewed computed tomography angiography and ventilation-perfusion scans. GV, SR and NN collected thrombus tissue from operating room. GV and IC (Ian Cummings) performed single cell RNA sample preparation, paraffin embedding and OCT cryopreservation. HFK performed scRNAseq computational work and HFK, SR, YRY and DC analyzed scRNAseq data. GV performed and analyzed *in vitro* experiments. NN performed CTEPH cell isolation and cell proliferation. IC (Issac Choi) performed tissue sectioning and slide preparation. GV, IC (Issac Choi) performed and analyzed immuno-histo chemistry and immuno-fluorescence imaging. AW performed and analyzed Circo plots of ligand:receptor interactions. All authors reviewed the results and approved the final version of the manuscript.

## Acknowledgements

We thank the Duke Human Vaccine Institute Genetics Analysis Core for preparation of scRNAseq libraries, the AML Laboratories for HE and MT staining, and Yasheng Gao in the Duke Light Microscopy Core Facility for help with live cell imaging and confocal immuno-fluorescence imaging. SR was funded by a Duke University Chancellor’s Discovery Award and an American Heart Association Transformational Project 19TPA34880033 Award. GV was funded by a Mandel Foundation Fellow award.

