## Supplemental data for "Single-Cell Analysis Reveals Distinct Immune and Smooth Muscle Cell Populations that Contribute to Chronic Thromboembolic Pulmonary Hypertension"

**Supplementary Table 1.** Clinical characteristics of subjects included in the study.

| Gender, % male | 60% |
| --- | --- |
| Race, (% white, % black) | 40%, 60% |
| Age, y, median [range] | 57 [45-63] |
| Weight, kg, media [range] | 91.4 [76.9-132.5] |
| NYHA class, % I/II/III/IV | 0 / 0 / 80% / 20% |
| Associated conditions |  |
| CAD | 60% |
| HTN | 80% |
| DM2 | 20% |
| CKD | 20% |
| Right heart catheterization |  |
| RA, mmHg, median [range] | 9 [4-20] |
| Mean PA pressures, mmHg, median [range] | 42 [40-64] |
| PVR, WU, median [range] | 7.7 [5.8-18.3] |
| Cardiac Index, L*min-1*m-2, median [range] | 1.8 [1.6 – 2.2] |
| Scintigraphy or CT angiography, % abnormal | 100% |
| Medical therapy, % (n) | 40% |
| Phosphodiesterase type V inhibitor | 40% |
| Soluble Guanylate Cyclase Inhibitor | 0% |
| Endothelin receptor antagonist | 20% |
| Prostacyclin analog | 20% |
| Combination therapy | 20% |

**Supplemental Figure 1.** Computed tomography angiography (CTA) and ventilation-perfusion (VQ) scan from three of the five subjects included in this study.

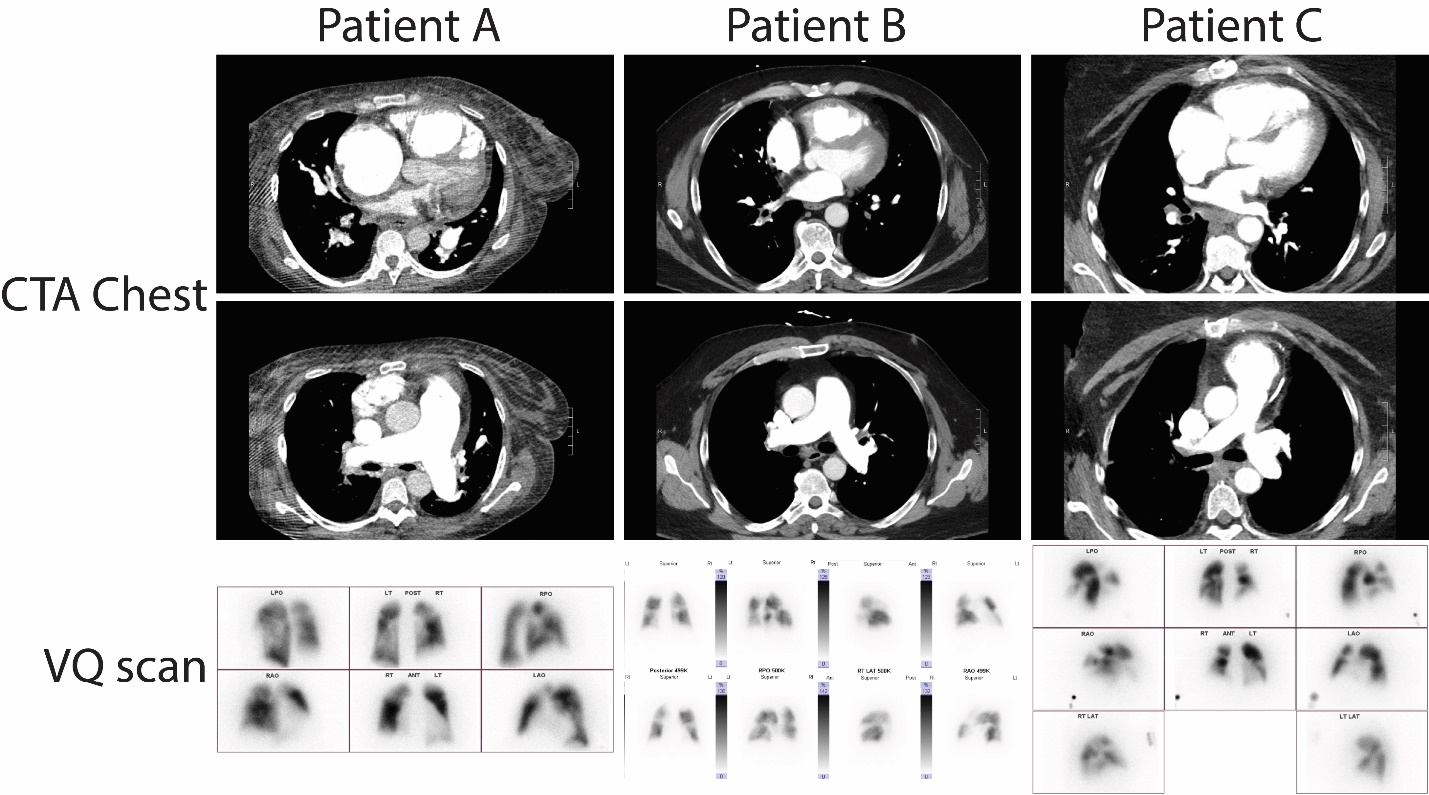

**Supplemental Figure 2.** Pictures of thrombus removed at the time of PTE surgery. Parts of the distal (segmental) thrombus were used for scRNAseq studies.

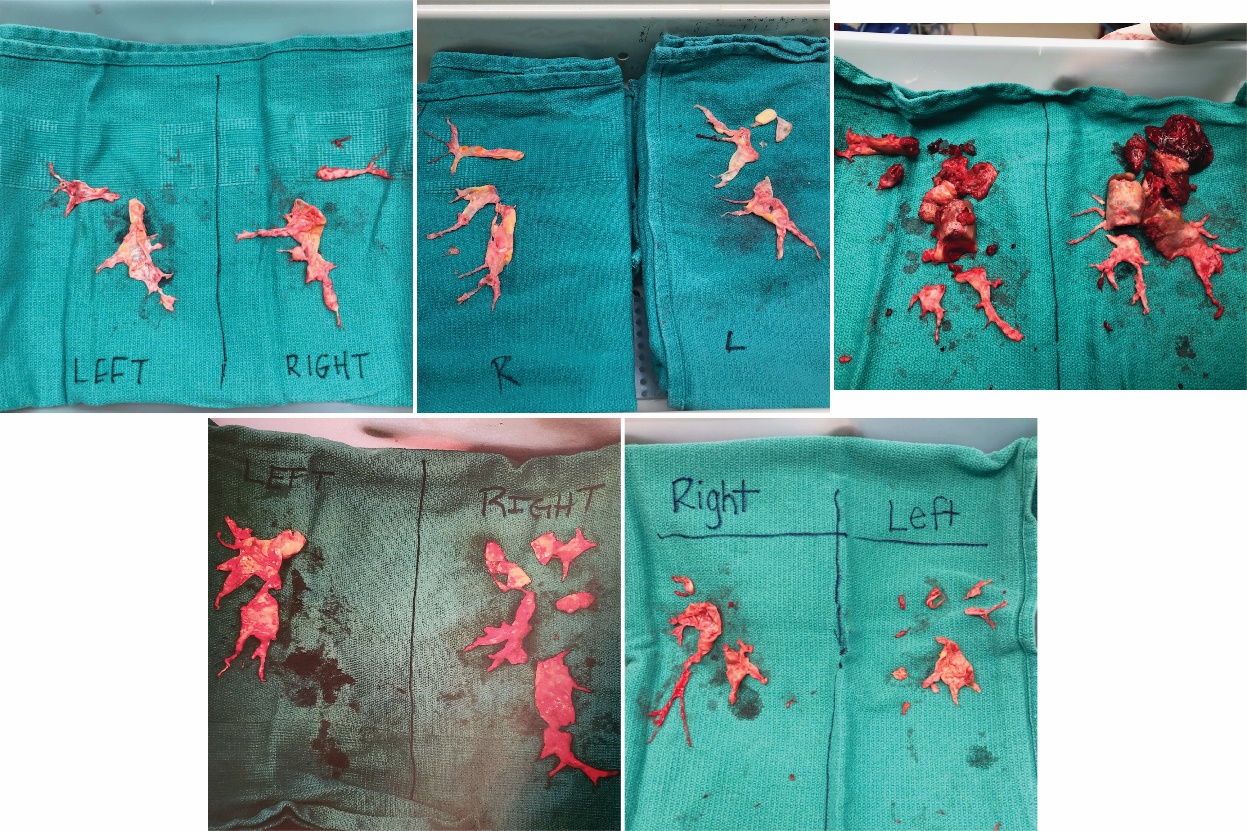

**Supplementary Figure 3.** Representative immunofluorescence images of macrophage, T cell, vWF and α-sm actin expression in CTEPH thrombus from three biological replicates.

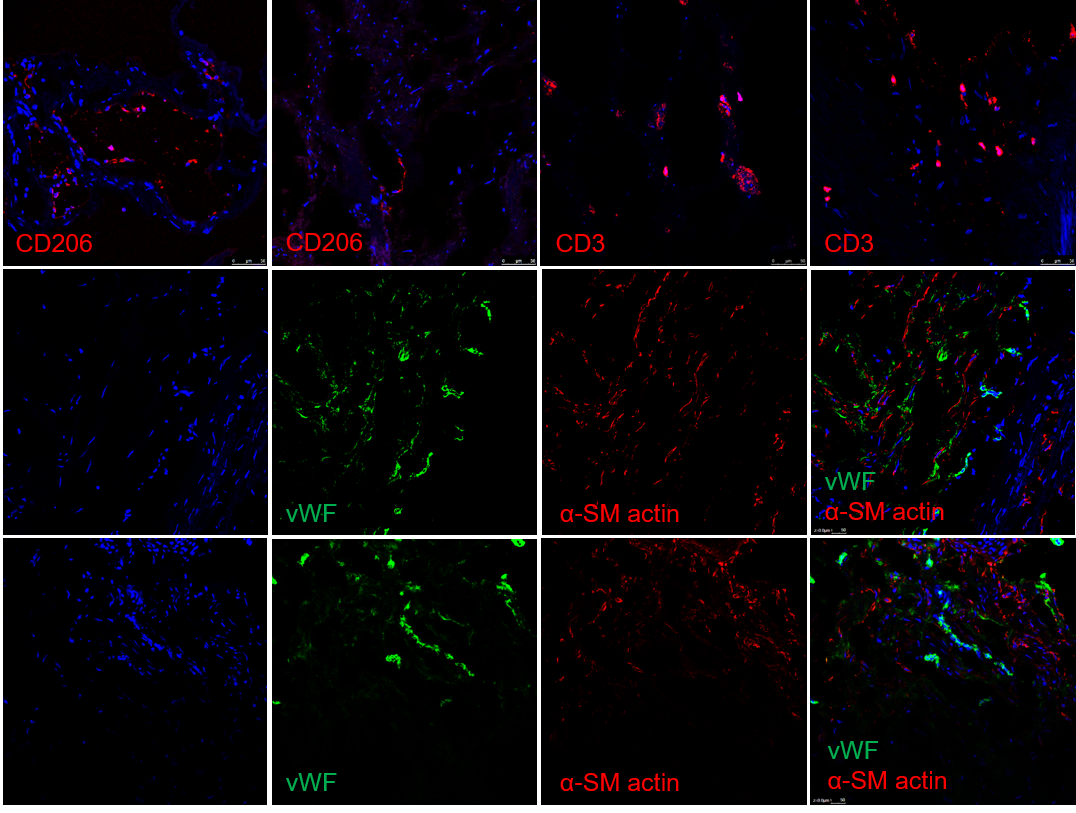

**Supplementary Figure 4.** Correlation of the largest five macrophage clusters with canonical macrophage markers. Mac-1 expressed highest levels of CXCR4; Mac-2: MerTK; Mac-3: CD11b, CD88/C5AR1, Mac4: CD68, CD163, CD206/Mrc1, CD169/Siglec1, CSF1R, Cx3cr1 and Lyve1; Mac-5: CCR2 and CD14, CD80, and CD86.

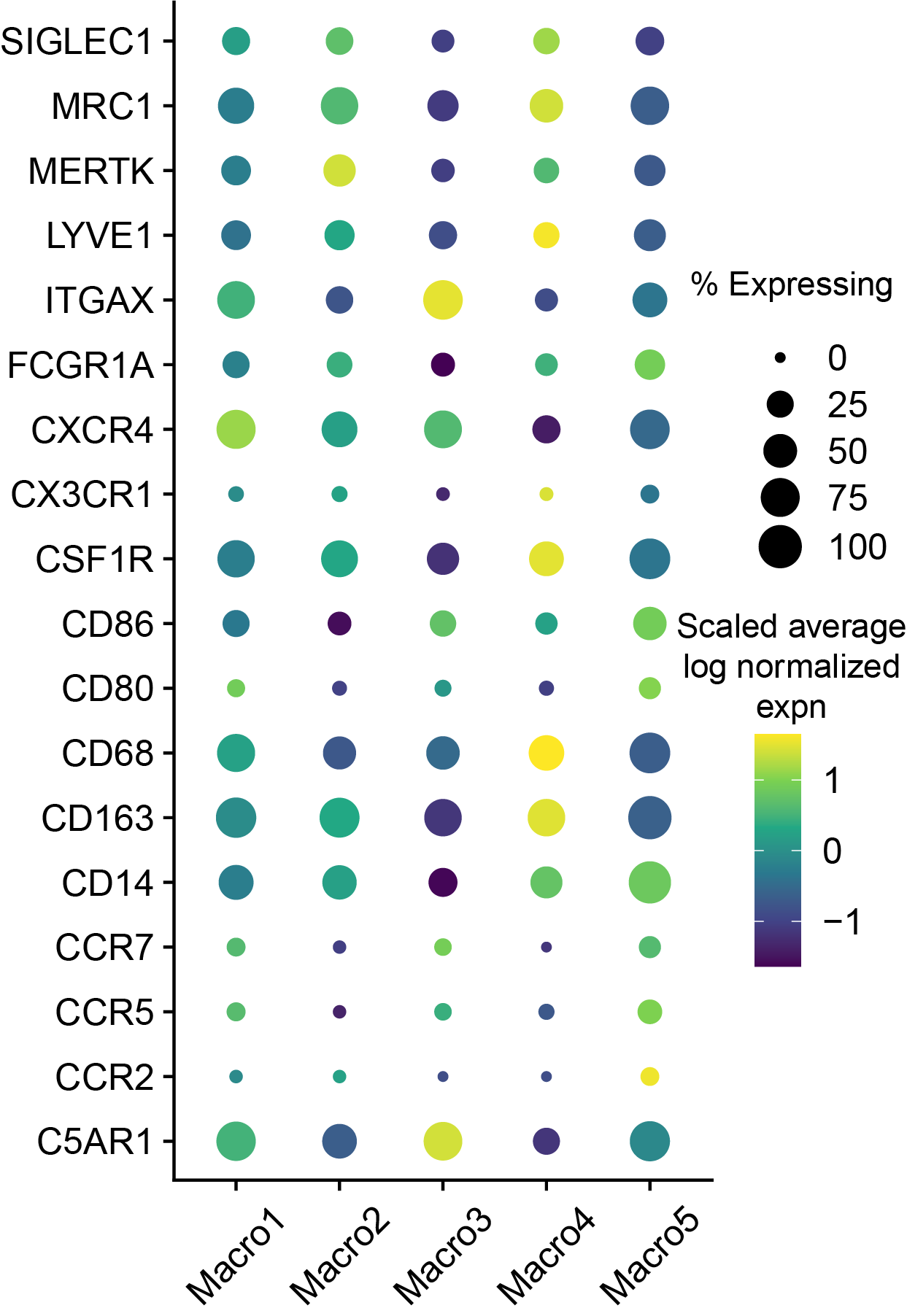

**Supplementary Figure 5.** Additional representative images of immunofluorescence of ECs, SMCs and myofibroblasts from three biological replicates.

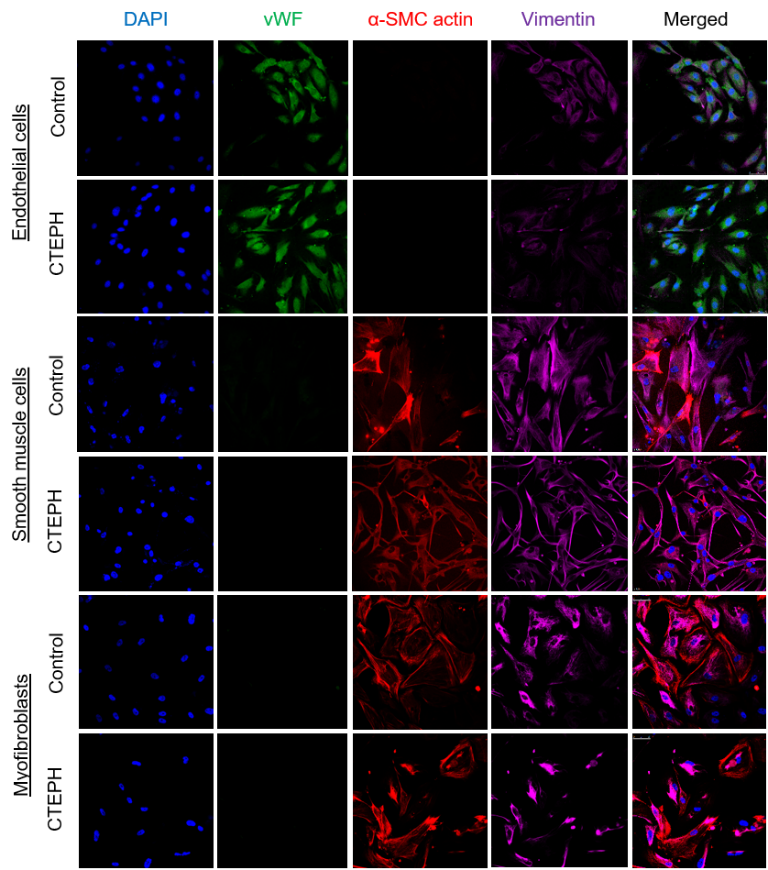

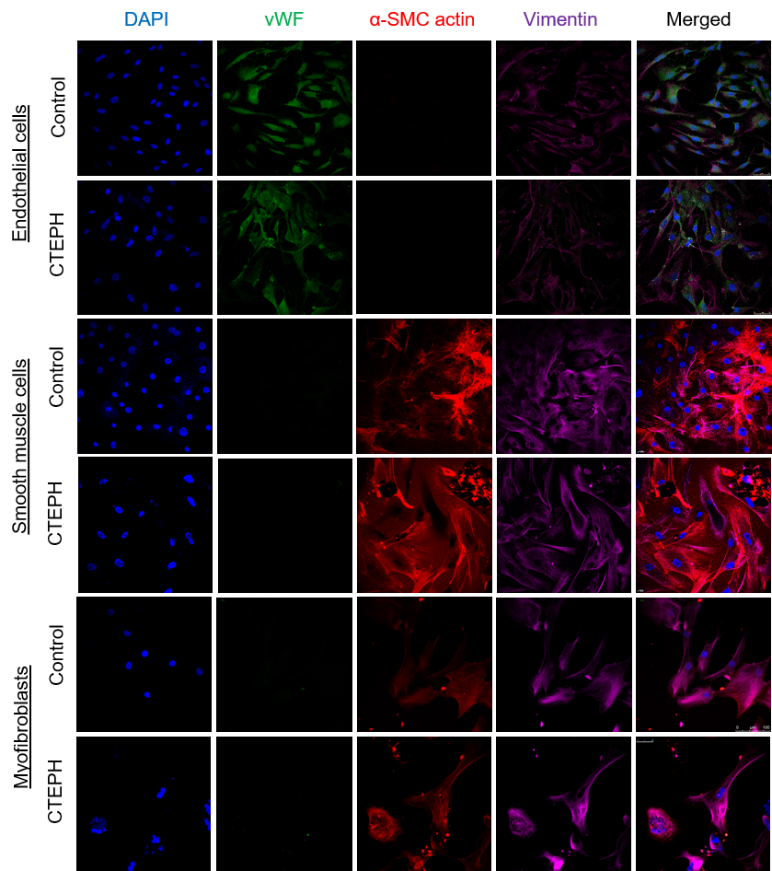

**Supplementary Figure 6.** Circo plots of ligand:receptor interactions between T cells, macrophages and smooth muscle cell clusters (data is presented as Ligands from specific cell type: Receptors from specific cell type).

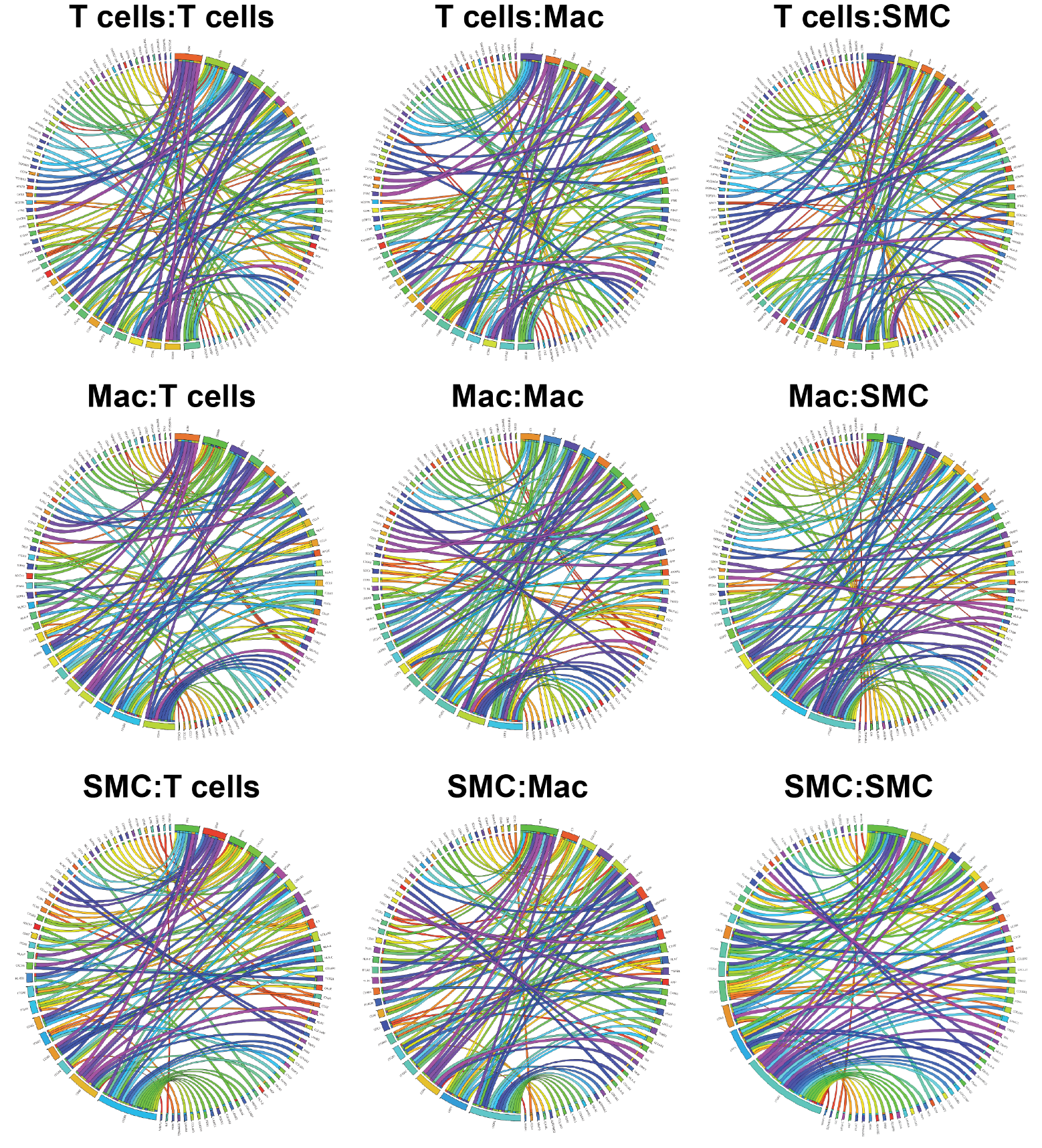

**Supplementary Figure 7.** Representative images of cultured CTEPH (**A**) SMC and (**B**) myofibroblast migration in response to thrombin and vorapaxar treatment.

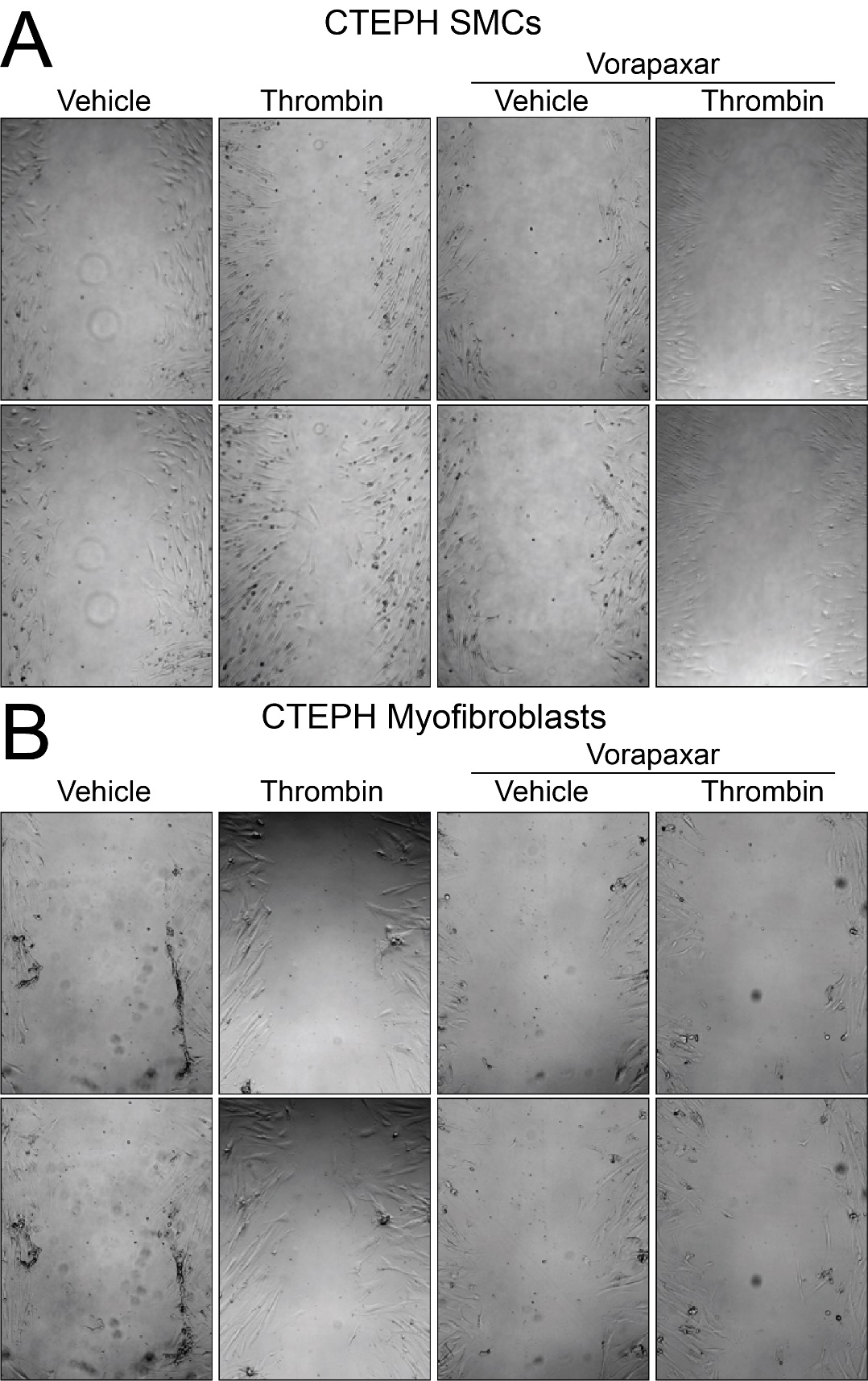
